# The Risk of Gulf Birds’ Functional Diversity Loss with Climate Change Uncovered Using Deep Learning Population Models

**DOI:** 10.64898/2026.03.19.712781

**Authors:** Liying Li, Junwen Bai, Shoukun Sun, Marcos Zuzuaggrei, Zhe Wang

**Affiliations:** National Center for Ecological Analysis and Synthesis, University of California, Santa Barbara; Santa Barbara, California, United States; Cornell University; Ithaca, New York, United States; University of Idaho; Idaho Falls, Idaho, United States; School of Engineering, University of California, Merced, Merced, California, United States

## Abstract

Climate change and sea-level rise (SLR) pose increasing threats to coastal ecosystems and biodiversity in the Gulf of America. Most efforts to anticipate these threats focus on species counts or range shifts, while changes in species functional diversity remain uncovered. We estimated the impacts of climate change and sea-level rise on hundreds of bird species’ populations and corresponding shifts in functional diversity. We used the generative deep learning method, Variational Gaussian Mixture Autoencoder (GMVAE), and Trait Probability Density analysis to study such impacts. We found that a generative GMVAE model uncovered species’ unobserved ranges, and that climate change reduced coastal ecosystem resilience and caused biodiversity loss across multiple dimensions, including functional richness, redundancy, evenness, and divergence. Surprisingly, the most impacted areas are not the exposed shoreline but the landward coastal transition zones. Specifically, shoreline functional diversity turned out to increase with climate change and sea level rise, whereas uplands showed declining functional diversity and increasing redundancy, indicating contraction of functional trait space. Furthermore, avian biodiversity expanded in coastal protected areas, serving as refugia embedded in a surrounding landscape where unique combinations of species traits are lost.

## 1. Introduction

Climate change and sea-level rises are fundamentally reshaping coastal geomorphology, altering ecosystem structure and biodiversity. Projections from the IPCC and U.S. federal assessments indicate substantial increases in chronic high-tide flooding and storm-driven coastal inundation over the coming decades, compressing intertidal habitats through a process known as coastal squeeze in the Gulf of Mexico (NOAA, 2022). At the same time, ocean warming, acidification, deoxygenation, and intensifying storm regimes compound these pressures across coastal habitats. Because many coastal birds rely on tidal flats, marshes, and shorelines for nesting, foraging, and migration stopovers, the loss or transformation of these habitats threatens their population persistence. These climate-driven changes, combined with growing human pressures on ecosystems (Olden et al., 2008), pose mounting threats to biodiversity. Land use change, fisheries bycatch for seabirds, pollution, and prey depletion drove widespread population declines and reductions in functional diversity across bird communities (Dias et al., 2019; Rosenberg et al., 2019; Saintilan et al., 2014). Quantifying the spatial patterns of these population and functional diversity changes is therefore critical for informing conservation strategies under accelerating coastal changes.

Species distribution models (SDMs) provide a practical framework for predicting species’ responses to environmental change (Botella et al., 2018; Elith & Leathwick, 2009) and can be applied to assess the impacts of climate change on Gulf birds. Recent advances in remote sensing, machine learning, and citizen science have improved predictive performance by capturing heterogeneous responses and by integrating a variety of data sources (Adadi, 2021). However, rare or difficult-to-identify species are underrepresented in conventional machine learning models due to data scarcity (Botella et al., 2018).

Embedding-based deep learning has enabled modeling a large number of species jointly, so rare species can be included in the modeling (Chen et al., 2018; Bai et al., 2020). These methods simultaneously normalize and project auxiliary information (environmental variables and coordinates) and vector-space embeddings (multiple species checklists) into a shared high-dimensional joint embedding space. Metrics such as cosine similarity (Chen et al., 2018) are used to measure the proximity of a focal species embedding to embeddings of other species and environmental variables, and to predict the focal species’ class (presence or absence). This is a promising approach for approximating biotic and abiotic relations for species distribution modeling.

The idea of borrowing parameters from common-species models to improve predictions for rare species has been also used in CORAL (Ovaskainen et al., 2025), a Bayesian transfer learning model (Suder et al., 2023). Both embedding approach and CORAL reflect a central idea in many joint species distribution modeling (JSDM) approaches (Norberg et al., 2019). These models, however, are typically trained within a fixed data domain. When domain shifts occur under climate change—such as the emergence of novel environmental conditions, species range shifts, or the introduction of new species—the learned species–environment relationships may no longer hold, potentially limiting predictive reliability (Miao et al., 2023).

Generative models are particularly well-suited for assessing biodiversity responses to climate change when domain shifts occur between training and prediction environments. Variational autoencoders (VAEs) learn latent representations of species–environment relationships, allowing models to generalize beyond the conditions observed in the training data (Kingma & Welling, 2022). Leveraging this property, Dinnage (2024) applied a generative VAE framework for zero-shot prediction (Mehta & Harchaoui, 2025), demonstrating accurate predictions for rare species and biodiversity monitoring regions with limited observations. Similarly, transfer learning approaches based on VAE models have been used to predict plant community composition in data-scarce regions by transferring knowledge from species distribution models trained in data-rich regions (Hirn et al., 2022). Because generative VAEs model the joint distribution of species and environmental variables through a shared latent space, they can maintain robust predictive performance even when applied to datasets characterized by different environmental conditions and species assemblages.

Despite these advances, significant gaps remain. First, class imbalance, often manifested as zero inflation, has not been explicitly addressed in these deep-learning species distribution models (SDMs), even though rare species and absence-dominated observations are pervasive in biodiversity datasets (Johnston et al., 2021). Second, most existing deep-learning SDMs focus primarily on species occurrence, which limits our ability to assess extinction risk and population responses to environmental and anthropogenic change (Thuiller et al., 2004). While occurrence predictions can identify potential habitat suitability, they provide limited insight into changes in species abundance and community structure. Third, biodiversity assessments that rely solely on taxonomic units as correlates of community richness are insufficient for evaluating ecosystem health (Larson et al., 2021). Species’ functional trait diversity is the mechanism behind biological responses to ecosystem change (Baker et al., 2021). Functional traits are morphological, physiological, or behavioral characteristics of organisms that can be quantified at various organizational levels, ranging from individuals to ecosystems. In communities where functional trait redundancy is low—or where functionally unique species are lost—ecosystem functioning can decline disproportionately (Biggs et al., 2020; Violle et al., 2017). Integrating functional trait information into biodiversity analysis, therefore, represents a critical direction for improving conservation prioritization and predicting biodiversity resilience.

To address these gaps, we applied a generative probabilistic deep-learning hurdle modeling framework to predict the abundance of 350 bird species across the Northern Gulf of Mexico under climate change and sea-level scenarios. The framework, previously developed by Kong et al., (2020). combines a Generative Gaussian Mixture Autoencoder with a hurdle likelihood to jointly model species coexistence and species population in a shared environmental space across large multispecies communities. This approach explicitly addresses the pervasive problem of zero inflation in biodiversity observations by separating occurrence and positive-abundance processes. Transfer learning and a contrastive loss function (Khosla et al., 2021a; Oord et al., 2019) enable the model to align the two separated stages of the hurdle model, each representing the alignment between species co-occurrences and species population correlation (Edmondson et al., 2021). Using this framework, we generated spatially explicit predictions of habitat suitability, species abundance, and community structure for Gulf birds with and without climate change and sea level rise impacts. We further integrated population-weighted functional trait metrics (Carmona et al., 2019) to link predicted taxonomic units to ecosystem functioning. This integration allows us to assess how environmental change affects not only species distributions but also functional diversity and ecosystem resilience.

We found an unexpected spatial pattern of biodiversity vulnerability. Contrary to expectations, the greatest biodiversity loss occurs in landward lower-upland zones rather than along exposed shorelines to sea level rises. Along shorelines, functional diversity increases as opportunistic species capitalize on episodic inundation and newly available habitats. In contrast, upland areas experience declining functional diversity and increasing redundancy, indicating a contraction of functional trait space driven by climate-induced habitat compression. We further identify protected reserves with appropriate nutrient management as potential functional refugia capable of buffering biodiversity loss. These fine-scale maps of species abundance and functional diversity shifts provide actionable insights for conservation planning and reserve management under with a changing environment.

## 2. Materials and Methods

### 2.1. Study area and data

Analyses were conducted at 3kmx3km spatial resolution on the northern Gulf of Mexico coast, including the southeastern parishes of Louisiana, the coastal counties of Mississippi and Alabama, and the western counties of the Florida panhandle. The study area aligned with the National Centers for Coastal Ocean Science project on Predicting Impacts of Sea Level Rise in the Northern Gulf of Mexico (NCOOS, 2017). All the required simulated data to create SLR scenarios are available for this study area, such as Bathymetry DEM (NOAA National Geophysical Data Center, 2010) and marsh productivity change with sea level rise for three National Estuarine Environmental Research Reserves: Apalachicola, FL, Weeks Bay, AL, and Grand Bay(Alizad et al., 2016). Our study drew on findings from studies examining the impact of SLR on coasts and coastal habitats, translating them into species response information to inform conservation and ecosystem management.

We used observation records from eBird (Sullivan et al., 2009) from 2019 to 2021, which included up to 115,575 checklist occurrences and counts after filtering, along with sampling-event metadata (e.g., protocol, duration, distance traveled, number of observers, start time). The filtering procedure followed the one described in (Li et al., 2025). Data was acquired and merged into one year for data augmentation. Checklists of all species were summarized into a table of species observations, with each shared locality ID represented as a single row in the table. The data version used in this study was the eBird Reference Dataset from 2025. To derive detections and non-detections, we restricted analyses to Complete Checklists—checklists where observers report all species they could identify during the sampling event—so that species not reported can be treated as absences on that checklist. Species responses were aggregated to the modeling grid with a 1km buffer. We modeled all 350 bird species (trimmed from 404 species observed by at least 10 observation each species) within the study area for two targets per 3km*3km grid cell: (i) occurrence (presence/absence) and (ii) abundance/count (non-negative), only excluding species with fewer than 10 observations over the three years through January, 2019 to December, 2021. Taxonomy and nomenclature follow the eBird taxonomy 2020.

Environmental covariates were assembled, tabulated, and standardized prior to modeling. Predictors included: topography (Amatulli et al., 2018)—mean elevation, elevation variability, slope, and aspect decomposed to northness and eastness; Land cover—proportional cover from C-CAP/land-cover classes aggregated to the grid (NOAA, 2024); climate-mean temperature and temperature range, and primary productivity of ecosystem- biomass concentration, and chlorophyll concentration. A full description of the list of data used and data sources can be found in this Supplementary Information. A climate change and sea level rise scenario is created by modifying the historical prediction surface with replaced temperature under climate change from Bio-ORACLE (Assis et al., 2024) and WorldClim2 (Fick & Jijmans, 2017) for ocean and land, respectively, and land cover classes with marsh migration (Marcy et al., 2011) (accessed at https://coast.noaa.gov/slrdata/). The projected temperature was acquired for CMIP6 SSP5-8.5, matching the intermediate high sea level rise scenario (1.2 m above the mean sea surface level by 2100) (Parris et al., 2012). We utilized Mean High Water and Salt Marsh Productivity data from the NOAA National Centers for Environmental Information (Alizad et al., 2018), which were modeled using Hydro-MEM (Alizad et al., 2016), to modify the historical bathymetry, digital elevation, and primary productive predictors (biomass concentration and chlorophyll concentration) for the three National Estuarine Research Reserves.

### 2.2. Generative Variational Autoencoder with Gaussian Mixture

Utilizing citizen science data for species occurrence and count observations requires a robust model structure to account for temporal and spatial data noise (Elith et al., 2006). AI SDMs have been proven to outperform traditional machine learning models to handle such noise (Lee et al., 2022). We follow a generative framework of Gaussian Mixture Variational Autoencoders (Bai et al., 2021a), which allows for zero-shot learning (predicting for species not seen during training) (Yan et al., 2024).

Limited observing efforts, constrained by resources supporting ecosystem monitoring, have significantly constrained our ability to define long-term and large-scale ecosystem and biodiversity patterns, especially at the species level. A location at a period of time that lacks strategic observation is not identical to a lack of biodiversity; however, if we quantify biodiversity based on biased observation, we will have a biased view of biodiversity. Zero-shot learning and prediction can identify a species niche even when a species only has a few occurrences in the training data.

The architecture consists of two parallel encoders, feature and label encoders, a shared latent representation, and two decoders, each designed to align and reconstruct feature and label modalities (Figure 1).

**Figure 1.**
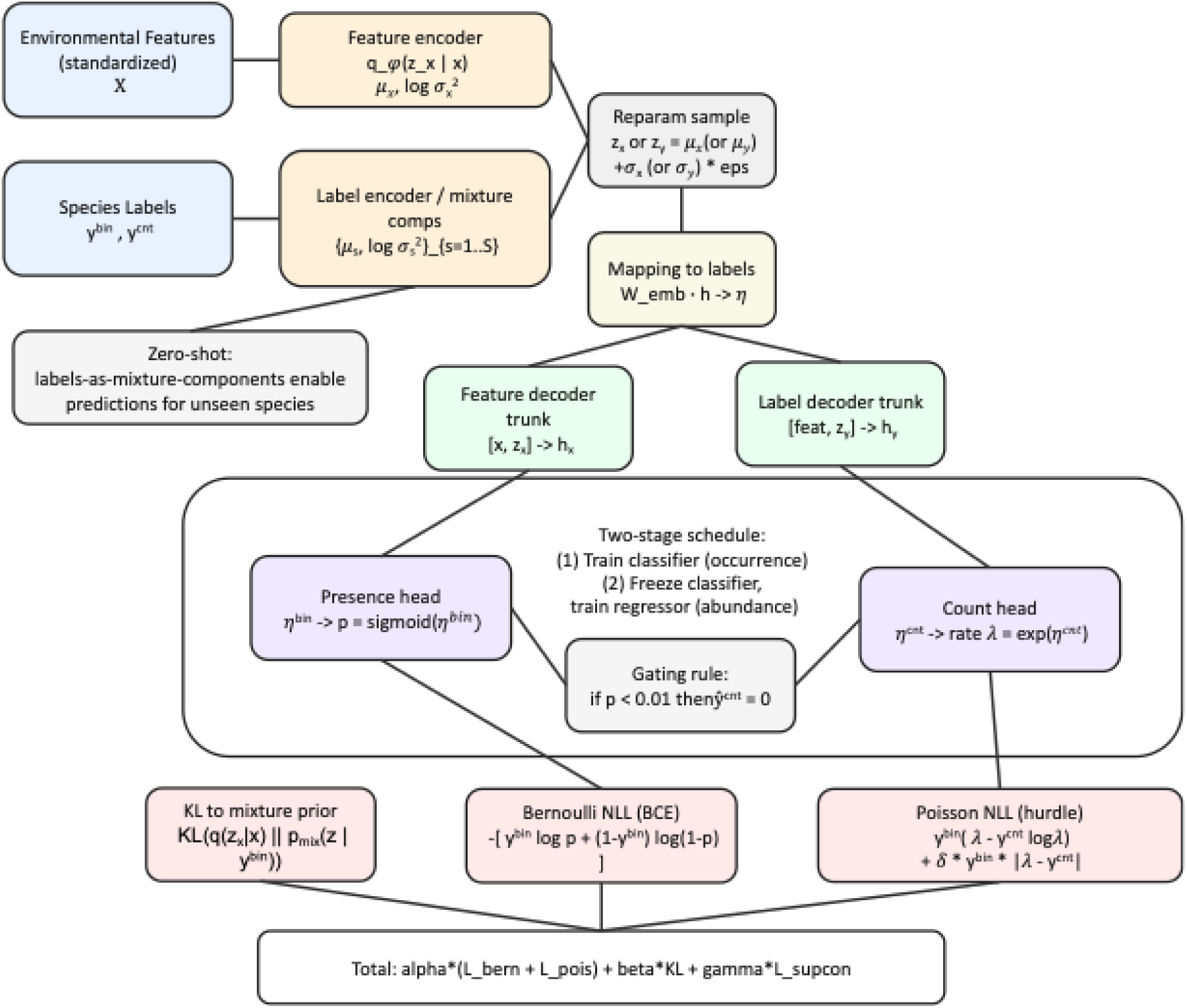
Diagram of the Generative Variational Autoencoder with Gaussian Mixture in a hurdle framework (gate rule, conditional Poisson loss, and two-stage training schedules) for species abundance predictions gated by occurrence predictions.

Environmental feature vectors Χ at a location/time (e.g., land-cover fractions, elevation stats, temperature means/ranges, chlorophyll, etc.) were first standardized and then passed through a multi-layer perceptron (MLP) (Rumelhart et al., 1986) (three fully connected layers of 256, 512, 256 neurons respectively) with rectified linear unit (ReLU) activations and dropout regularization. Outputs µ_x_ (mean), log a ^2^ (log variance) are mapped to a latent Gaussian distribution (ɛ∼N(0, *I*)) and reparametrized by µ_X_ and log σ^2^, generating a reparametrized latent sample z_X_ for the feature label path. Species occurrence or abundance labels j were embedded and projected through a symmetric encoder consisting of fully connected layers (embedding dimension 512 and 256 neurons) with ReLU activations and dropout. The resulting output (mean µ_s_, variance log a ^2^ per species) was parameterized into a Gaussian distribution. Reparameterization step (Equation 1) that ensured differentiability while capturing uncertainty in the latent variables:

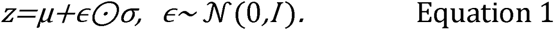

And each label was further modeled as a component of a Gaussian mixture, enabling mixture-based latent modeling. The Gaussian Mixture Variational Autoencoder (GMVAE) prior (Bai et al., 2021) in the latent space allows each species label to be associated with a distinct mixture component, supporting zero-shot prediction (Yan et al., 2024).

Both encoders produced distributions from which latent codes were sampled using the reparameterization trick (Equation 1). Latent variables were passed through two decoding pathways: The Feature Decoder h_y_ concatenated latent codes with input features and projected them through two fully connected layers to generate normalized embeddings in the shared space. The Label Decoder h_x_ concatenated latent codes with feature embeddings and projected them through fully connected layers with leaky ReLU activation to generate label embeddings. A reconstruction branch directly attempted to reconstruct the original features from the latent codes, ensuring information preservation. Both feature and label decoders were aligned into a shared embedding space. A learned linear transformation then mapped embeddings back into the label space (W_emb · h -> 77), enabling final predictions of species occurrence probability or abundance.

Each VAE uses the same internal components as described above (feature encoder/decoder and label encoder/decoder) but is trained with task-specific likelihoods inside a composite objective. Let L_NLL_ denote the prediction negative log-likelihood terms (both the label-decoder path and two feature-decoder-based–based paths), L_KL_ the latent KL divergence (GMVAE regularization) (Kingma & Welling, 2022), and L_CPC_ the contrastive alignment loss (Bai et al., 2021b; Khosla et al., 2021). Loss contributions were weighted such that NLL received primary emphasis (α=10), KL divergence was scaled by β=6 to balance reconstruction and regularization, and CPC alignment was scaled by γ=1 (Kong et al., 2020). During training, the weighted sum of loss (Equation 2) was minimized:

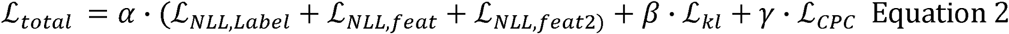

### 2.3. Two-stage hurdle structure

A two-stage hurdle approach (Kong et al., 2020) was used to model species abundance distribution for 350 species informed by occurrence modeling results. A hurdle model is a two-part model that specifies two separate processes, one for generating zero values and another for generating values given that they are non-zero, for handling count observations with excess zeros (Edmondson et al., 2021)

In this study, the first stage is using the Multivariate Probit Model (Chen et al., 2016) to model the presence/absence of each species, capturing interspecies and environment–species covariance. (Bai et al., 2020). This stage is treated as a foundation model (Dinnage, 2024) that provides ecological context. Building on the occurrence distribution, a second fine-tuned regression abundance model was trained using species count data with rich context informed by the occurrence model. For classification, the label decoder projects into a probability space through a sigmoid output layer, yielding predicted occurrence probabilities for each species (Presence head: 77bin -> p = sigmoid(77^bin)). For regression, the decoder projects to a continuous output layer with a linear activation, yielding predicted species abundances.

Because both tasks share the same latent embedding, gradients from classification and regression jointly shape the latent space (Count head: 77_cnt_ ➔ rate A = exp(77_CTit_)). This encourages the model to learn representations that are predictive of both species presence/absence and relative abundance, rather than optimizing them in isolation; therefore, it was named as cross-task regulation. This integration allows the model to flexibly shift between predicting occurrence and abundance depending on available data, while maintaining a unified ecological embedding space.

The NLL loss uses a hurdle formulation with two probabilistic heads: a Bernoulli component for presence and a Poisson component for conditional counts (Bishop, 2016; Huffer et al., 2008). During the occurrence (classification) phase, the classifier outputs a logit η_is_^(bin)^, converted to a probability p_is_=σ(η_is_^(bin)^) with the logistic sigmoid σ (⋅). The Bernoulli negative log-likelihood (equivalently, binary cross-entropy), averaged over samples and species, is

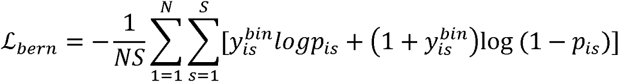

Let x_i_ denote environmental features for sample i=1,…,N and y ^bin^∈{0,1} the presence/absence of species s=1,…,S. When present, the observed count is y ^cnt^∈ N.

During the Abundance (regression) phase, the standard Poisson log-likelihood with log link (Cameron & Trivedi, 2013; Kingma & Welling, 2022) was used, which was restricted to the positive (post-hurdle). The regressor outputs an unconstrained value η_is_^(cnt)^ that parameterizes the Poisson rate λ_is_=exp (η_is_^(cnt)^). A small L_aux_ auxiliary term to stabilize rate estimates. Counts are modeled only when a species is present via the hurdle mask m_is_=y_is_^bin^. The presence indication masked the Poisson loss to respect the hurdle structure.

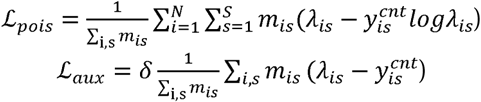

### 2.4. Species range and population outputs

The deep hurdle model was used to predict species habitat suitability and population across the whole study area using the prepared prediction surfaces for different scenarios. Prediction scenarios include historical and climate change scenarios. To acquire species range, raw sigmoid outputs from the classifier’s feature path were binarized using data-adaptive, species-specific thresholds: for species with batch mean probability > 0.01, we used the batch column mean as the threshold (probability > mean → 1), and for lower-prevalence species we used a fixed threshold of 0.05 (probability > 0.05 → 1). The classifier’s probabilities are used to gate abundance predictions as a hurdle during regressor training. If the presence probability is less than 0.01, the predicted abundance is set to 0.

Bias is more prominent in count observations than occurrence observations. That is, observers may record thousands of birds without counting them; therefore, the larger the count, the greater the error may be. To mitigate observer-related biases, we train the abundance model using log-transformed counts rather than raw counts. Predictions are then back-transformed to the original scale and rounded to integers for evaluation with mean squared error (MSE). The alignment between where species are present and the predicted number of individuals is assured by the Contrastive Loss (Bai et al., 2021). The predicted population is not hindered by the predicted accuracy because arbitrary occurrence thresholds were used, as described above. Instead, abundance values are expected abundance derived by multiplying the predicted occurrence probability by the predicted counts.

### 2.5. Model training and evaluation

Model training was conducted using Pytorch Lightning (Falcon, 2019) with GPU acceleration where available. The Adam optimizer was applied with separate initial learning rates of 5e^-3^ and 2e^-4^ for classification and regression, respectively. These rates decayed by a factor of 0.5 if the validation loss did not improve for four epochs, using a StepLR scheduler. All experiments were repeated across 70%/15%/15% train/validation/test partitions using a fixed random seed (42) to ensure robustness, and results were reported as the mean ± standard deviation of evaluation metrics. TorchMetrics was used for accuracy, RMSE evaluation. Models were trained for up to 500 epochs each for classification and regression. The regressor VAE was trained while the classifier was frozen/evaluation. During this phase, the regressor consumed the classifier’s sigmoid presence probabilities (computed without gradient) to gate abundance predictions. No gradient clipping and no early stopping were applied. The mini-batch size was 4096 for all experiments.

Classification and regression were evaluated using different sets of metrics. Classification used Accuracy, F1-score, Precision, Recall, and Intersection over Union (IoU). These metrics captured different aspects of predictive performance, with IoU providing a robust measure of overlap between predicted and observed species presence. Additionally, reliability diagrams and Brier scores were used to assess the degree to which predicted probability models matched observed frequencies. Regression used Root Mean Squared Error (RMSE) and Mean Absolute Error (MAE). RMSE emphasized large deviations between predicted and observed abundance values, while MAE provided a more interpretable measure of average prediction error. Together, these metrics provided a comprehensive evaluation of predictive accuracy, robustness, and ecological interpretability across the two modeling tasks.

### 2.6. Trait Density Probability Analysis

Species traits were sourced from AVONET (Tobias et al., 2022), a global compilation of avian morphology and ecology with species-level summaries derived from >90,000 measured individuals across 11,000+ species. However, only species-level mean traits were provided in AVONET. Intraspecific trait variation arising from genetic differences and phenotypic plasticity can be large enough to alter interaction outcomes and community assembly. Therefore, consideration of stochasticity in intraspecific trait differences is necessary to mitigate the risks of biasing trait distribution change predictions due to environmental changes.

We used continuous trait summaries in AVONET as standardized means for species-level continuous traits and categorical traits as categorical assignments. Continuous traits were converted to pseudo-individual values by parametric jittering with independent Gaussian draws using the species-level means from AVONET. A conservative coefficient of variation (5%) was imposed to reflect unobserved within-species variability in the absence of trait-specific variance estimates from AVONET. To align traits with our observation table, we expanded each species’ trait record to the number of observations for that species (n ). When the trait was missing or zero, values were left as NA. Categorical traits (e.g., trophic niche, habitat, migratory status) were replicated deterministically across the n rows. This strategy preserves species identity while yielding an observation-aligned trait matrix in which continuous traits incorporate modest within-species dispersion to accommodate intraspecific variation that affects community assembly (Bolnick et al., 2011) when only species means are available.

We quantified species’ functional structure using the Trait Probability Density (TPD) approach implemented in the R package TPD (Adams et al., 2025; Carmona et al., 2019). All trait analyses were conducted in the R programming language (R core team, 2021). The TPD framework represents each ecological unit (species, community) by a probability density function in trait space, enabling overlap-based measures of richness, dissimilarity, redundancy, and divergence.

### 2.7. Defining trait spaces

Continuous traits (and, where mixed traits were present, ordination scores from) were scaled to zero mean and unit variance before analysis. Species-level probability densities were estimated using multivariate kernel density estimation, and returns species/population TPDs were evaluated on a regular grid. We set the probability isopleth at alpha = 0.95 to define the high-probability support (non-zero region) and controlled grid resolution via two divisions. We didn’t use a Gower–PCoA reduction to analyze the mixed traits space of continuous traits and categorical traits. Instead, we categorize species by their categorical traits and built three pair-wise 2-dimensional trait spaces, namely Beak Length/Beak Depth (indicating foraging trait space), Tail length/Hand Wing Index (indicating movement trait space), and Tarsus Length/Habitat Density (Indicating social behavior space). Tarsus length represents species size, and we regard species of different sizes as having distinct social behavior, as many studies have suggested (Adams et al., 2025; Delestrade, 2001; Weeks et al., 2020).

From species to communities, community-level densities were obtained for each of the three spaces, which mixes species probability density, weighted by their relative abundances in each sampling unit (here we used historical and sea level scenarios). This produces the Trait Probability Density (TPD) for each scenario, which is directly comparable for evaluating species resilience in response to sea level rise. We summarized and reported the functional structure using richness, evenness, redundancy, and divergence, all of which were computed on the probabilistic trait space.

### 2.8. Functional diversity indicators

Functional evenness and divergence are two distinct components of functional diversity compared to functional richness and redundancy. The richness and redundancy indicators aim to estimate the amount of niche space filled by the community and the overlapping of these niche spaces. These two aspects of functional diversity do not take into account the species abundance distribution, giving equal weight to rare and common traits (Karadimou et al., 2016). In order to take into account that rare traits may play a more significant role in the ecosystem functioning of a community, we consider all four indicators in our study to assess climate change impacts on community compositions.

Functional evenness and divergence focus on the distribution of individuals in this trait space (Lechêne et al., 2018). Functional evenness represents the uniformity of species abundance across occupied trait space; high values indicate similar contributions of variety strategies, whereas low values indicate dominance by a few strategies and gaps in trait use. Functional divergence measures how far species with high abundances at a location are from the center of the community’s functional space; a higher value indicates a greater prevalence of extreme traits.

If richness increases, evenness can either increase or decrease. When it increases, it means new niches or habitats become available to species with disturbance and dynamic inundation gradients, and newly introduced functional groups can coexist with prior functional groups. No single strategy dominates among species. On the contrary, if evenness decreases, it means most of the population is concentrated in a few strategies with functional imbalance for rare traits. If richness decreases, evenness may remain because a community could lose a specialized species (lowering functional richness), but the remaining species might still have very similar abundances, which keeps functional evenness high. Alternatively, the total functional space could shrink, but one species could become highly dominant, lowering functional evenness.

## 3. Results

### 3.1. Species population decline with SLR

We projected species populations (Figure 2) with climate change, modified coastal inundation levels, and marsh migration trajectories. The generative deep hurdle model produced abundance maps for bird species, comparing the historical baseline to the climate change scenario along the northern Gulf Coast. Across species, coherent coastal bands in the historical maps break into patchier inland clusters under SLR, especially along major waterways—indicating displacement rather than uniform decline in population. The scenario maps showed persistence clustering along tidal creek networks and near estuary mouths, while back-barrier/impounded cells lose suitability—i.e., places with better tidal exchange fare better. The hotspot of species distribution geography becomes more fragmented as a result of adaptation to sea level rise.

**Figure 2.**
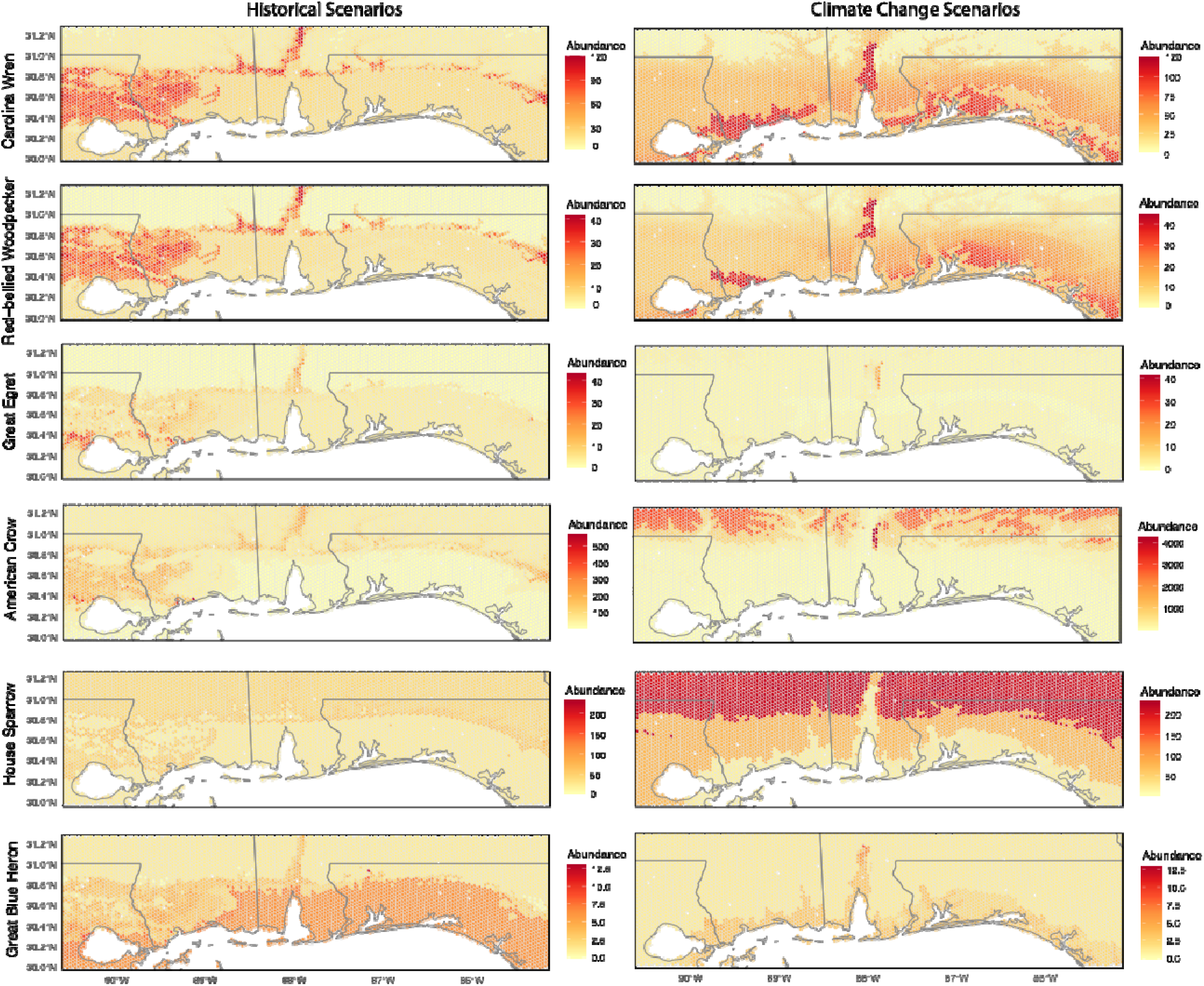
Maps of expected abundance change from historical scenarios to climate change scenarios.

The highest losses occur where inland migration is blocked by roads/levees, while sites connected to channels retain or gain suitability. The climate change scenario maps showed persistence clustering along tidal creek networks and near estuarine mouths, while back-barrier/impounded locations lost suitability (i.e., places with better tidal exchange fared better). Zooming to wetland specialists, a contract near the coast and a shift up-estuary were observed. For example, high-abundance cells of Great Egret and Great Blue Heron, located along the immediate shoreline, weaken under SLR, with hotspots reappearing along river mouths and in higher-elevation marsh and back-bay zones inland. This pattern is consistent with tidal-marsh “coastal squeeze,” where inundated lowlands are not fully replaced inland.

Forest birds, such as the Carolina Wren and Red-bellied Woodpecker, show a modest coastal retreat but retain strong holdouts in the interior. They observed a reduction in abundance in low-lying coastal forests under SLR, while interior upland/river-corridor cells remained stable or intensified. The signal suggests their sensitivity to flood-driven habitat change at the coastal fringe, but resilience persists where canopy structure remains intact inland. American Crow and House Sparrow are generalists displaying the smallest scenario deltas. Abundance remains comparatively stable, with slight inland shifts tracking developed corridors and higher ground. A diffuse interior presence, with subtle redistribution offsets, mitigates any local coastal dips in species abundance. In sum, local coastal losses redistribute abundance inland and up-estuary for wetland birds, cause minor edge retreats for forest birds, and leave generalists essentially unchanged aside from subtle inland shifts. The losses at the lowest elevations are compensated by persistence inland abundance, which is consistent with landward habitat migration constrained by topography and development.

### 3.2. Uncovering the hidden patterns of functional richness

Through abundance-weighted trait analysis, we found that local functional richness is, historically, highest in a narrow coastal ribbon that tracks the Mississippi Sound barrier-island chain, the Grand Bay marsh complex, and the Mobile Bay–Tensaw Delta. Eastward, elevated values continue along back-barrier and estuarine shorelines of Perdido–Pensacola Bay, Santa Rosa Sound, and Choctawhatchee Bay (Figure 3). These hotspots coincide with shallow, structurally diverse wetland mosaics that host many flight-strategy and dispersal types (e.g., strong fliers, short-hop residents, wide-ranging coastal movers). Richness declines both offshore (in open Gulf cells south of the islands) and inland (toward the northern edge of the frame), both of which lack the fine-grained wetland complexity that supports diverse assemblages. These areas occupy the cooler color bins on the historical map (roughly the 30–40 range for movement trait richness).

**Figure 3.**
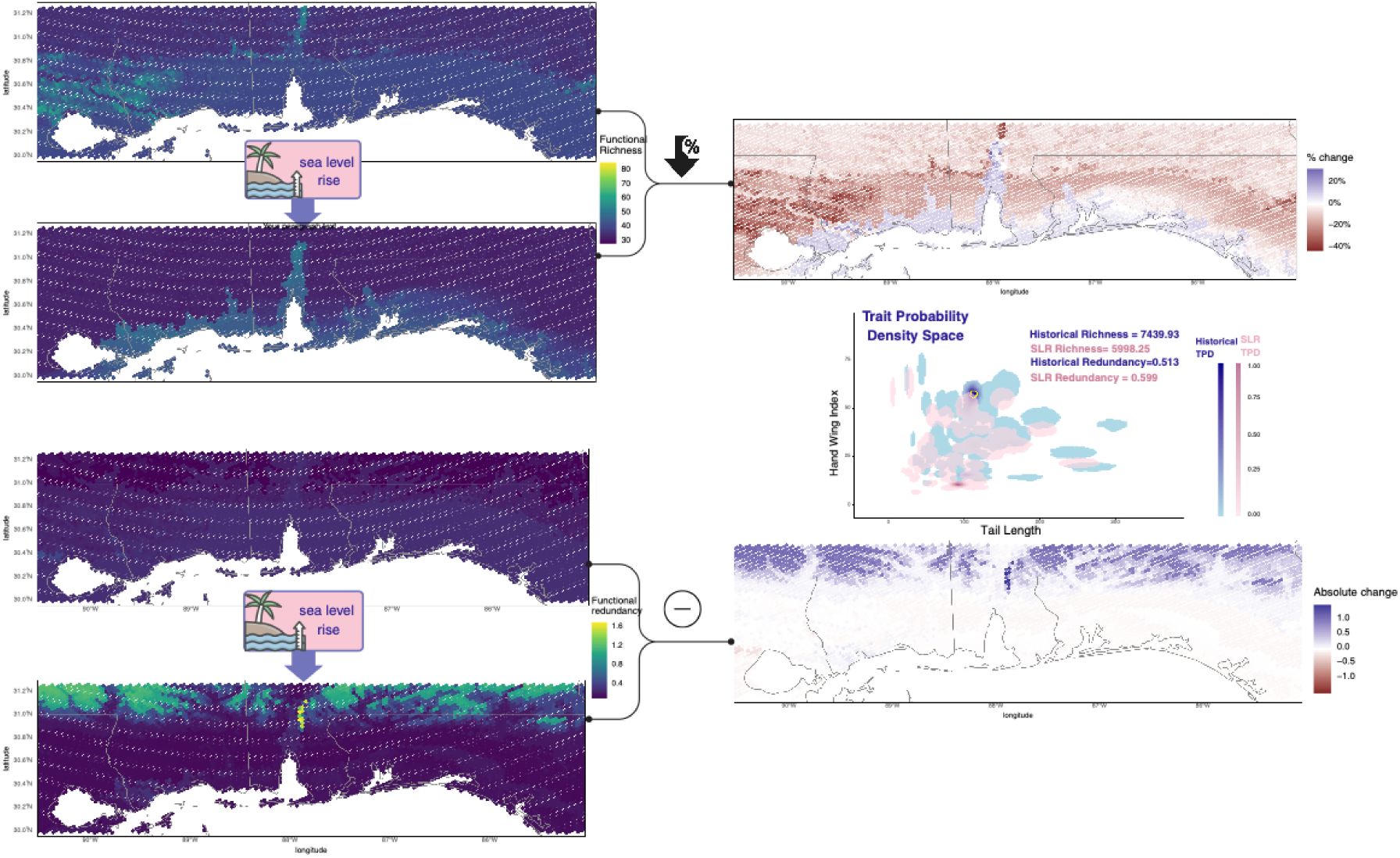
**Abundance-weighted functional richness and redundancy of 350 species movement traits with and without sea level rise impacts, and their changes with sea level rises, accompanied by trait probabilistic hypervolumes** (density probability of the full range of species traits combination). Color scales represent either trait diversity index values or trait diversity change values, the Hand Wing Index, and the Tail Length composite movement trait space. If the trait index values are pretty small, we use the absolute change value to prevent extremely large change values

Under climate change and SLR, the overall palette shifts cooler. Coastal cells closest to the shoreline show localized increases in functional richness while maintaining steady redundancy. These gains likely reflect short-range redistribution into newly wetted edge habitats and the persistence of tidal corridors that continue to support diverse movement modes. The largest richness declines form a coast-parallel band inland of the shoreline — roughly between the back-barrier edge and lower uplands (−20% to −40% in the percentage change panel). Hotspots persist most clearly in sheltered bay areas and a few back-barrier reaches, but they are narrower and less intense than historically, reflecting the loss of low-elevation marsh and supratidal habitats that supported a wide spread of movement strategies. Moving slightly inland and uplands, changes are modestly negative. In short, SLR drives coastal thinning and segmentation of movement functional richness.

High redundancy means ecosystems can absorb species losses while still maintaining key functions. Functional redundancy is highest along the same coastal mosaic where habitats and resources are broad and repeated—Mississippi Sound’s marsh–barrier system (Cat/Ship/Horn/Petit Bois–Dauphin Island), the Mobile Bay–Tensaw Delta complex, and the western Florida Panhandle embayments from Perdido–Pensacola through Santa Rosa and Choctawhatchee—reflecting multiple species filling similar roles within productive, shallow estuarine settings with SLR. Redundancy is comparatively lower offshore (open water Gulf cells) and in inland/upland cells where fewer species co-occur and trait overlap is limited (Figure 4). Under the sea-level-rise scenario, Movement functional redundancy is relatively stable along the immediate coast, remaining moderate through much of the barrier-island/estuarine corridor. By contrast, a broad inland/back-barrier belt exhibits a marked increase in redundancy. From a regional standpoint, redundancy increases slightly overall (from 0.513 to 0.599), meaning regional trait assemblages become more compositionally concentrated as shorelines retreat.

**Figure 4.**
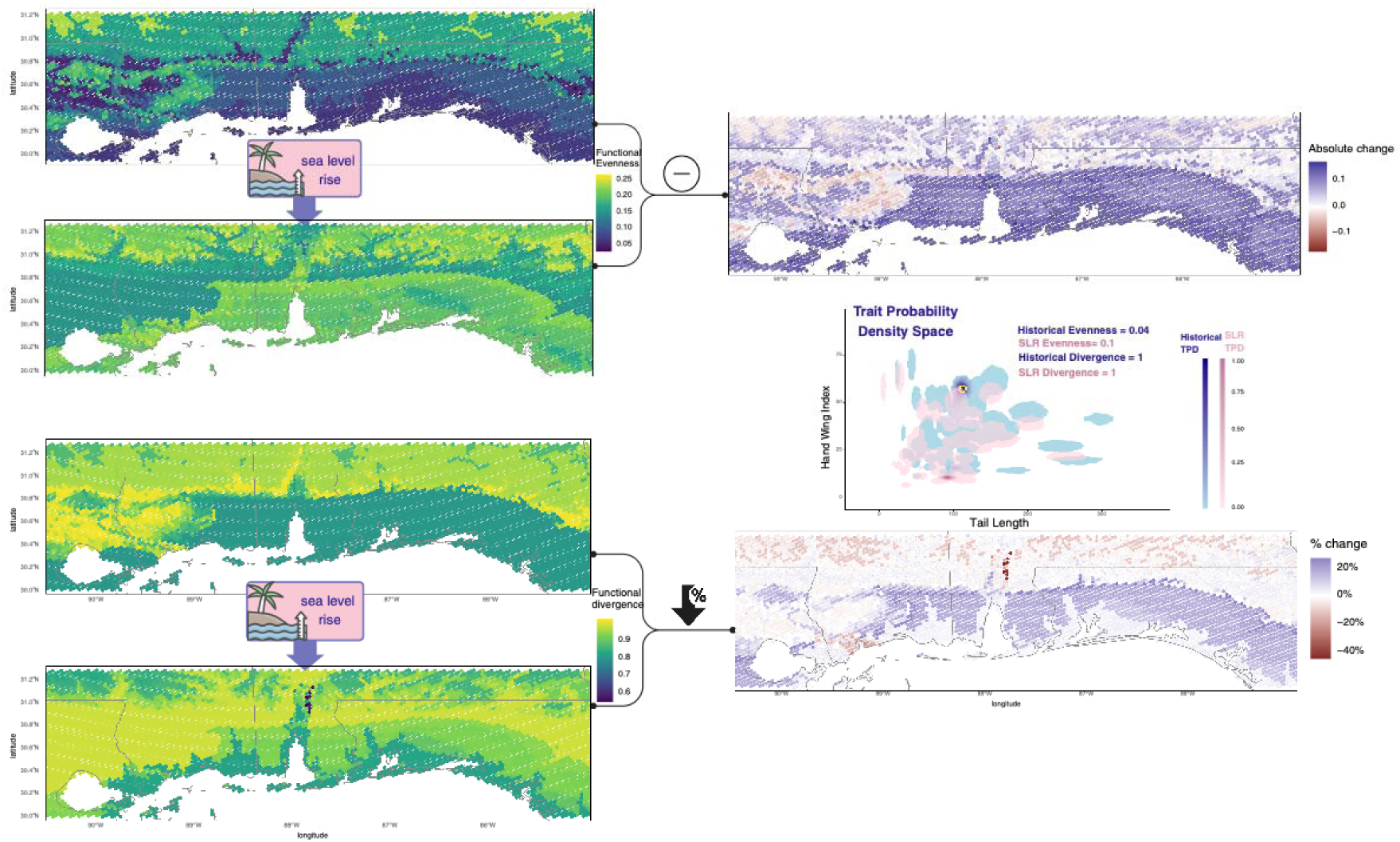
Abundance-weighted functional evenness and divergence of 350 species movement traits (Hand wing index and tail length) with and without sea level rise impacts, and their changes with sea level rises, accompanied by trait probabilistic hypervolumes (density probability of the full range of species traits combinations). Color scales represent either trait diversity index values or trait diversity change values, the Hand Wing Index, and the Tail Length composite movement trait space. If the trait index values are pretty small, we use the absolute change value to prevent extremely large changes.

The contracting richness and rising redundancy inland prove functional homogenization and biodiversity loss with SLR. Surprisingly, redundancy increases most significantly in uplands, meaning that upland species loss is characterized by a smaller number of species, but with distinctive movement strategies lost, and the remaining species cluster into overlapping roles. The directly exposed coastal cells didn’t see either a decrease in richness or an increase in redundancy. While even where inland richness losses are modest, the sharp redundancy gains signal reduced ecosystem versatility and lower resilience to additional stressors (e.g., storms, heat waves, prey shifts), as fewer complementary movement modes remain to sustain processes when conditions change.

These results suggest a shift from diverse, complementary coastal assemblages to more redundant inland assemblages under SLR from a regional perspective. Management actions that conserve or restore trait-diverse coastal wetland mosaics (e.g., living shorelines, sediment/nourishment to maintain back-barrier marsh platforms, protection of estuarine corridors) should help retain movement strategy breadth where it is currently highest, while efforts inland should prioritize maintaining complementarity (not just species counts) to avoid further erosion of functional versatility.

### 3.3. Uncover the pattern of functional evenness

In Figure 4, by contrast, high divergence may indicate communities that provide unique functions but are fragile if species disappear. A community where traits are evenly represented is more resilient to environmental fluctuations.

Evenness is spatially structured and shifted into a fragmented pattern with climate change. Historically, Evenness is lowest along the immediate coast (dark tones represents 0.05–0.10), especially around the Mississippi Sound barrier–island rim and the margins of Mobile Bay and the western Florida Panhandle embayments. These cells are dominated by a few movement strategies (e.g., shoreline specialists), yielding an uneven, spread functional trait space (in the trait density space). Higher evenness (0.15–0.25) appears in more interior cells, where assemblages contain a more balanced mix of movement types.

Under the SLR scenario, evenness increased broadly and became more spatially continuous. The shoreline corridor shifted to the dominance of a few coastal strategies. The largest increases in functional evenness occur along the Mississippi Sound corridor, spanning Bay St. Louis and the Grand Bay marsh complex, and extending across the. Mississippi and Alabama barrier island chain.

The absolute-change map confirms that SLR flattens the distribution of movement strategies across space (from 0.04 to 0.1) and the removal of rare functional traits: evenness rises almost everywhere, with the shoreline catching up but still lagging inland, consistent with a regional trend toward more uniform use of trait space uplands. This indicates markedly more even partitioning of movement functions among the species that adapted to SLR.

Historical pattern of functional divergence is highest inland and along the back-barrier/northern tier (yellow, 0.85–0.90), and lower along the immediate shoreline and open Gulf cells (teal, 0.70–0.80). The strongest inland values form an arc from coastal Mississippi into Alabama and the western Florida Panhandle, while coastal margins of Mississippi Sound, Mobile Bay, and the Perdido–Pensacola / Santa Rosa–Choctawhatchee shorelines remain comparatively lower. Under SLR, functional divergence shifts seaward: a coast-parallel belt from Mississippi Sound through Mobile Bay to the western Florida embayments increases into the higher classes, producing a more continuous yellow swath along the back-barrier/shoreline corridor. Conversely, the interior/upland band shows declines (cooler greens/teals). The percentage change map confirms the widespread positive changes (blue, up to 20%) hugging the coast and negative changes (brown, −20% to −40%) inland, with a localized decrease hotspot north of Mobile Bay.

SLR favors edge-weighted (extreme trait values), more extreme movement strategies in cells near the coast (rising functional divergence), while inland assemblages become more center-weighted (falling divergence), consistent with the inland increases in redundancy and evenness that were observed. In practical terms, coastal communities trend toward specialization at trait extremes, whereas inland communities flatten toward more generalist mixes as habitats reorganize with SLR.

### 3.4. Peeking inside the space of species’ ecosystem functions

We quantified community-level trait-based metrics and generated figures of trait hypervolumes: richness (volume of hypervolumes), redundancy (overlap/backup among similar trait combinations), evenness (balance of use across the occupied space of the hypervolume), and divergence (edge- vs center-weighted use). The whole-region-scale community’s realized functional hypervolume—the portion of trait space occupied by species and their abundances —was presented in Figure 5. We found trait richness contracts under SLR, indicating loss of unique combinations, while evenness generally increases, i.e., the combinations that remain are used more uniformly. The directions of change in redundancy and divergence differ by trait group, revealing how the contraction unfolds.

**Fig 5.**
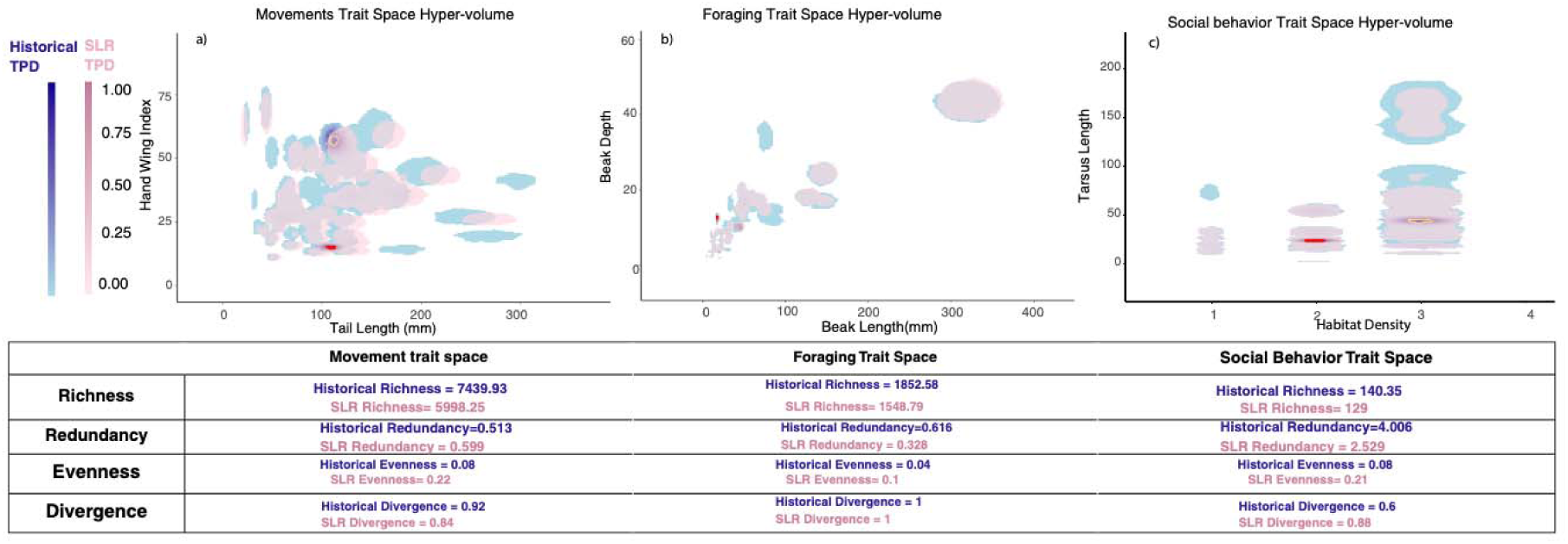
Functional trait probabilistic hypervolumes across the study area (probability density of the full range of species traits combinations of all present species) in three functional trait spaces with and without sea level rise impacts: movement, foraging, and social behavior trait spaces, and the table quantitative functional trait diversity index values. Function trait diversity index includes richness, redundancy, evenness, and divergence. Hand Wing Index and Tail Length combination composites movement space, Beak Depth and Beak Length composites foraging trait space, and Tarsus Length and Habitat Density composite social behavior space. The adjacent table lists all functional diversity indices for all three spaces in scenarios with and without sea level rise. Historical scenario without sea level rise is represented by blue color hypervolumes and index numbers, and sea level rise scenario is represented by pink color hypervolumes and index numbers. Highlight circles indicate the top 25% of the highest probability trait combinations within each space (yellow circle for historical scenario and red circle for climate change scenarios).

The movement hypervolume shrinks (richness 7,439.93 → 5,998.25), while redundancy rises (0.513 → 0.599) and evenness increases (0.04 → 0.10). Divergence stays high (≈1 → 1), implying that, despite volume loss, occupancy continues to weight the edges of movement space—extreme movement strategies persist—but with greater local overlap and a more even spread among those survivors.

The foraging hypervolume also contracts (richness 1,852.58 → 1,548.79), but here redundancy declines sharply (0.616 → 0.328), evenness rises (0.08 → 0.21), and divergence increases (0.60 → 0.88). Together, these shifts indicate pruning of intermediate strategies and a shift toward the edges of foraging space (e.g., more extreme bill/foraging morphologies), with less backup among similar strategies. The system becomes leaner and more specialized, not merely more overlapped.

Social behavior hypervolume shows a modest richness decline (140.35 → 129), a large drop in redundancy (4.006 → 2.529), a marked rise in evenness (0.08 → 0.22), and a decline in divergence (0.92 → 0.84). Abundance thus shifts toward the center of social-behavior space (fewer extreme social strategies), with less overlap/backup among similar social types but more even contributions of those that persist.

These complementary responses help reconcile spatial maps where richness declines while evenness (and sometimes redundancy) increases. Functionally, the community retains some extreme movement types, becomes more specialized in foraging, and less polarized in social strategies—all while overall trait breadth narrows. Management should therefore (i) protect trait-diverse coastal mosaics to maintain movement strategy breadth where it remains highest, (ii) buffer against foraging specialization risk by conserving prey/habitat diversity and cross-habitat linkages (given falling foraging redundancy), and (iii) sustain social-strategy complementarity (colonial vs solitary, territorial vs non-territorial) to avoid further center-ward collapse. Because redundancy changes are trait-group-specific—rising for movement but falling for foraging and social traits—resilience will depend on which functions are targeted: coastal connectivity may still be buffered by movement overlap, whereas trophic and social processes could be more fragile under continued sea-level rise.

## 4. Discussion and conclusion

Anticipating that biodiversity changes in response to global environmental change (Olden et al., 2008) to inform conservation and restoration actions requires ecological forecasts that resolve patterns at both species and community levels. Conservation has traditionally focused on species counts (richness, abundance, endemism), but functional traits and their community-level patterns add another crucial dimension for planning and resilience.

Our AI hurdle model—coupling occurrence and abundance prediction in a two-stage manner while aligning both predictions to a shared latent space—captured fine-scale, nonlinear species–environment relationships while borrowing statistical strength across taxa via learned species embeddings. This structure enabled generative (zero-shot) expected-abundance predictions for sparsely observed species by positioning them in the label-embedding space informed by the rich ecological context of species niche.

Spatially explicit functional diversity maps (richness, redundancy, evenness, divergence) revealed a consistent change gradient in response to climate change and sea-level rise (SLR). First, coastal cells (barrier-island rims, back-barrier marshes, estuarine margins) generally show increases or maintenance of functional richness with redundancy holding steady, while evenness rises, indicating more uniform participation among movement strategies without dominance by a few traits. Second, the largest richness losses concentrate in a coast-parallel, mid-elevation lower uplands just landward of the shoreline, where habitat conversion and edge losses exhibit the strongest decline of biodiversity. Third, upland areas exhibit strong increases in redundancy despite small declines in richness—an inland functional homogenization signal consistent with the consolidation of overlapping strategies as unique combinations and diversity are lost.

Trait-group-specific hypervolume analyses clarify how this reorganization occurs at the regional community level. The movement hypervolume shrinks regionally yet retains edge-weighted functional diversity and gains redundancy and evenness, implying fewer distinct movement trait combinations overall but more overlap and more uniform use among those that remain after SLR. The foraging hypervolume contracts as divergence increases and redundancy declines, suggesting pruning of intermediate strategies and a shift toward more extreme foraging modes with reduced resilience of functional traits. The social-behavior hypervolume narrows modestly, evenness rises, and divergence falls, indicating a center-ward shift with fewer extreme social strategies or traits.

When we studied the coastal reserves within the study area (Grand Bay NERR, MS; Weeks Bay NERR, AL), their joint hypervolumes expand or are maintained and fill more evenly across movement, foraging, and social traits; redundancy rises notably for movement and social domains, while foraging redundancy remains comparatively low. In contrast, at the whole-region scale (coast + inland), the overall hypervolume contracts, mid-elevation cells drive biodiversity loss, and uplands also experience trait homogenization. Together, these results position the reserves as functional diversity refugia—places where trait breadth and balanced use persist or improve under SLR—embedded in a surrounding landscape that is losing unique combinations and converging toward fewer traits.

Our analysis showed that no single biodiversity metric tells the whole story. Sea-level rise (SLR) often raises evenness—abundances become more uniformly distributed across the trait combinations that remain, which can be read as greater short-term resiliencies against further disturbance. But read in isolation, evenness can be misleading. Our trait diversity maps and hypervolumes show that these evenness gains frequently occur within a contracted trait space with lower biodiversity (functional diversity). In other words, communities may appear more “balanced” while simultaneously losing unique trait combinations that underpin rare or specialized functions, which can be understood as irreversible harm to ecosystems and biodiversity.

These spatially explicit findings can be translated into clear guidance for biodiversity conservation. 1) Trophic failure remains the weak link, as coastal foraging redundancy remains low despite gains in richness and evenness. 2) Strengthen prey and habitat diversity and cross-habitat linkages by restoring oyster–marsh–seagrass mosaics, securing fresh–brackish flow pulses (seasonal freshwater releases), plus local nutrient/turbidity management and seasonal protections for sensitive flats. 3) Preserve social-strategy complementarity by protecting both colonial and solitary sites and roost–forage corridors, supported by dynamic buffers and evidence-based predator management. 4) Convert coastal capacity into continuity by keeping tidal corridors open and edge habitats intact (living shorelines, marsh-platform maintenance, strategic sediment placement). 5) A dynamic buffer zone can be used around colonies to sustain functional traits and optimize management at reserve scales. 6) Finally, plan beyond reserve boundaries—use corridor easements and rolling setbacks to secure landward pathways and blunt inland homogenization.

Taken together, our results provide a tractable blueprint for biodiversity conservation actions. The approach used here to model the entire ecosystem of bird species demonstrates a novel way for ecologists and decision-makers to forecast and monitor biodiversity everywhere.

## Acknowledgments

We thank Professor Ben Halpern (Director, NCEAS) for insightful comments and guidance during manuscript development, and Courtney Scarborough (Deputy Director, NCEAS) for helpful comments and assistance with manuscript editing. We thank Dr.

Daniel Fink for meaningful discussions on the methodology. This paper is a result of research funded by the National Oceanic and Atmospheric Administration’s RESTORE Science Program (ROR - https://ror.org/0042xzm63) under award NA22NOS4510203 to the National Center for Ecological Analysis and Synthesis (NCEAS) at the University of California, Santa Barbara, and part of the Gulf Ecosystem Initiative.

## Author contributions

Conceptualization: LL, JB, MZ, ZW

Methodology: LL, JB, SS

Investigation: LL, SS, ZW

Visualization: LL

Supervision: LL Writing—original draft: LL

Writing—review & editing: LL, JB, MZ, SS, ZW

## Data and materials availability

All data are available in the main text or the supplementary materials

## Supplementary Materials

Materials and Methods

Supplementary Text

Figs. S1 to S6

Tabe. S1. Species abundance maps for additional bird species not listed in the Main Text-continued

References

